# Structural factors of RGS14 regulating hormone-dependent phosphate transport mediated by NPT2A

**DOI:** 10.1101/2024.03.29.586499

**Authors:** W. Bruce Sneddon, Suneela Ramineni, G. Emme Van Doorn, John R. Hepler, Peter A. Friedman

## Abstract

The NPT2A sodium phosphate cotransporter-2a mediates basal and parathyroid hormone (PTH)- and fibroblast growth factor-23 (FGF23)-regulated phosphate transport in kidney proximal tubule cells. Both basal and hormone-sensitive transport require sodium hydrogen exchanger regulatory factor-1 (NHERF1), a scaffold protein with 2 PDZ domains. NPT2A binds to PDZ1. RGS14 persistently represses hormone action by binding to PDZ2. RGS14 contains an RGS domain, two Ras/Rap-binding domains (RBD1, RBD2), a G protein regulatory (GPR) motif, and a carboxy-terminal PDZ ligand. The intrinsic RGS14 domains cannot explain its regulatory effects on hormone-sensitive phosphate transport because these actions are mediated not only by the PTH receptor, a G protein-coupled receptor (GPCR) but also by fibroblast growth factor receptor-1 (FGFR1), a receptor tyrosine kinase that is not governed by G protein activity. Here, we sought to identify the RGS14 structural elements mutually controlling PTH and FGF23 action. RGS14 truncation constructs lacking upstream sequence and the RGS domain were fully functional. Removing the linker sequence between RGS and RBD1 domains was sufficient to abrogate RGS14 action. Examination of the alpha-helical linker region suggested candidate serine residues that might facilitate regulatory activities. Ser^266^ and Ser^269^ replacement by Ala abolished RGS14 regulatory actions on hormone-sensitive phosphate transport and binding to NHERF1. The results establish that RGS14 actions require targeted phosphorylation within the linker and an intact PDZ ligand.

---

Parathyroid hormone (PTH) and fibroblast growth factor-23 (FGF23) regulate extracellular phosphate homeostasis by controlling sequestration of the NPT2A Na-phosphate cotransporter (SLC34A1) from kidney cell membranes. NPT2A internalization decreases phosphate absorption and promotes urinary excretion. The regulatory action of PTH (1,2) and FGF23 (3,4) on NPT2A requires the PDZ protein NHERF1 and, as we recently described, is tempered by Regulator of G protein signaling-14 (RGS14)^2^ (5). NHERF1 contains tandem PDZ domains. NPT2A binds PDZ1, whereas RGS14 binds the PDZ2 regulatory domain. RGS14 expression in human kidney cells stabilizes the [NPT2A:NHERF1] complex, blocking PTH and FGF23 actions.

RGS14 contains an RGS domain, tandem Ras/Rap-binding domains (R1/R2), and a G protein regulatory (GPR or GoLoco) motif (6) Activated Gα-GTP subunits bind the RGS domain, exerting GAP activity toward heterotrimeric G proteins to terminate GPCR signaling. Tandem RGS14 Ras/Rap-binding domains (RBDs, R1, R2) bind activated H-Ras-GTP R1 (7) and also interact with Rap2-GTP and Raf kinases, and Ca^2+^/CaM and CaM-dependent protein kinase (CaMKII) (8,9). The GPR motif binds inactive Gαi1/3-GDP, terminating guanine nucleotide exchange (*i*.*e*., GDI activity) and anchoring Rgs14 at the plasma membrane (10). Primates and ruminants express a 20-amino acid longer, full-length RGS14 protein terminating in a PDZ ligand that is absent in mice, rats, and most other species due to the presence of a UAG stop codon in exon 15 (11).

Human RGS14, containing a PDZ ligand (-DSAL), binds NHERF1, whereas rat Rgs14 lacking the PDZ ligand does not. Naturally occurring PDZ ligand variants disrupt RGS14 binding to NHERF1, thereby abrogating RGS14 actions (5). These findings established the role of RGS14 in controlling PTH- and FGF23-sensitive phosphate transport and underscored the requirement for the RGS14 PDZ ligand in this regulatory activity. The canonical RGS14 regulatory domains, associated with G proteins, cannot explain the effects on hormone-sensitive phosphate transport inasmuch as RGS14 controlled both PTH actions, mediated by its cognate GPCR and also those of FGF23 that are transduced by the structurally unrelated receptor tyrosine kinase FGFR1 (4,5). These observations suggested that the characterized RGS14 domains could not explain the elements in RGS14 governing its action. We hypothesized that RGS14 regulatory activity requires both an intact PDZ ligand and an upstream element. Based on these considerations, we analyzed the RGS14 structural determinants for mutual PTH and FGF23 action on phosphate transport mediated by NPT2A.

## Results

### Deletion mutants identify the RGS14 region responsible for regulating hormone-sensitive Pi uptake

RGS14 contains an RGS domain, tandem Ras/Rap-binding domains (R1/R2), and a G protein regulatory (GPR or GoLoco) motif (6,12) (Fig. 1A). We designed serial truncation mutants starting at the N-terminus to define the RGS14 structural determinants involved in PTH- and FGF23-sensitive phosphate transport. The described functional RGS, R1/R2, domains and the flanking interlinking regions were progressively deleted (Fig. 1A) to ascertain if they were required for RGS14 action on hormone-regulated phosphate transport. Truncation construct 1 removed the N-terminal sequence upstream of the RGS domain and retained the remaining protein. Deletion 2 cleaved the N-terminus and RGS domain, leaving the linker region, downstream R1/R2 domains, and GPR motif. Further shortening in construct 3 eliminated the linker, while construct 4 removed the two RBD domains, leaving the GPR motif and the C-terminus with its PDZ ligand. Truncation constructs 1-4 were transfected into HEK cells to test their expression. All were detectable by immunoblot, and each conformed to its expected size (Fig. 1B). WT-RGS14 was confirmed to be 63 kD. Truncation construct 1, which eliminated the N-terminus up to the RGS domain, ran at 56 kD. Truncation construct 2, in which the N-terminus and RGS domain were deleted, was detected at 43 kD. Truncation construct 3, which further deleted the linker between the RGS and R1 domains, ran at 30 kD. Truncation construct 4, which removed the N-terminus, RGS, R1, and R2 domains but left the GPR motif and PDZ ligand, was detected at 14 kD.

**Fig. 1.**
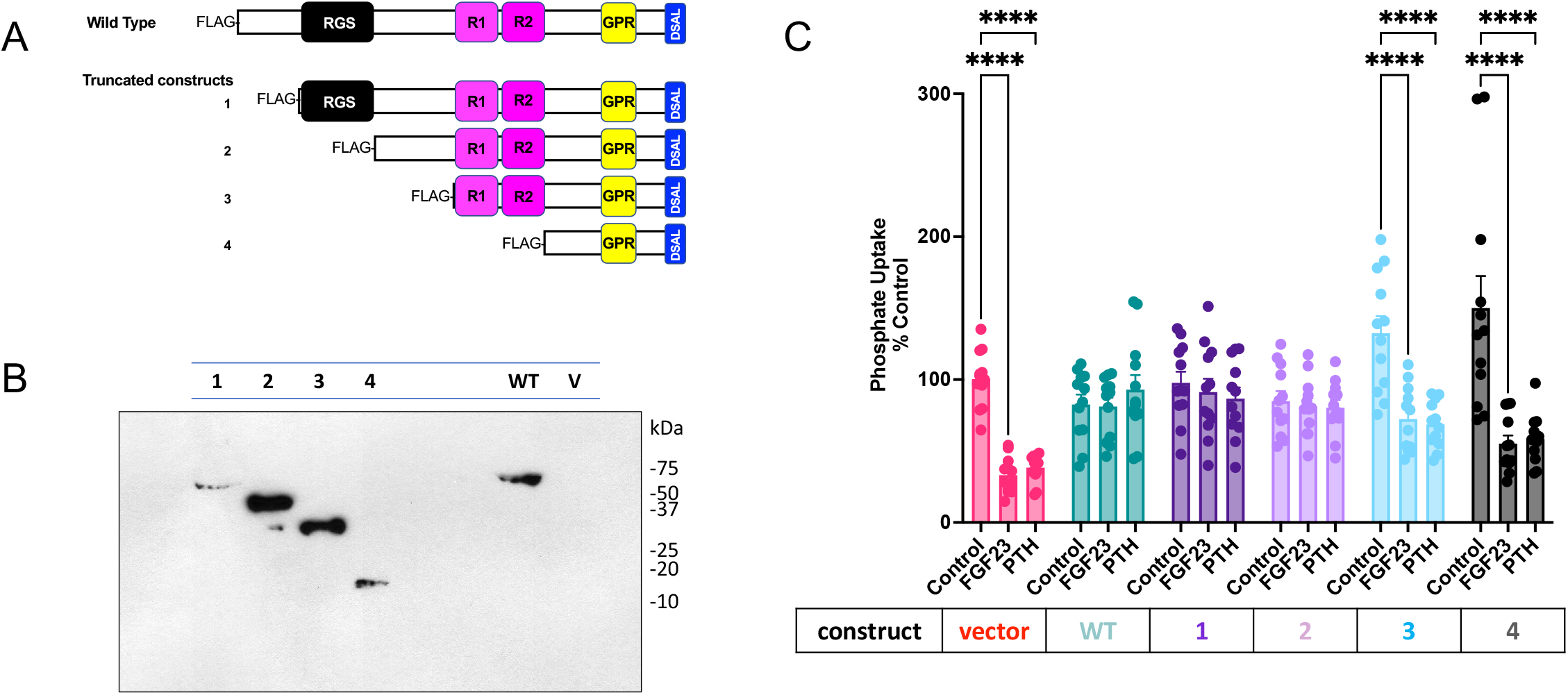
RGS14 blockade of hormone-sensitive phosphate uptake requires the linker region between the RGS and R1 domains. RGS14 truncation mutants were generated to identify functional domains suppressing hormone-regulated phosphate transport. *A*. Serial deletions from the N-terminus of RGS14 were generated, as outlined in Experimental Procedures. Previously characterized functional domains are depicted in the full-length (WT) RGS14 and the truncation mutants (1-4). Regulator of G protein Signaling (RGS) domain, Ras-binding domains 1 and 2 (R1, R2), G protein regulatory (Go-Loco) motif, and -DSAL PDZ ligand are depicted. *B*. Transfected WT-RGS14 and truncation mutants (constructs1-4) expressed in HEK293 cells. Molecular weight markers (kDa) are presented for reference. Empty vector-transfected cell lysate (V) served as a negative control. *C*. Hormone-sensitive phosphate uptake was measured in OK cells transfected with WT-RGS14 or the indicated truncation mutant. Phosphate transport was normalized to baseline phosphate uptake under control conditions (100%). Where indicated, cells were treated with 100nM FGF23 or PTH before phosphate uptake measurements. *n*=6. ****p<0.001 vs control.

Empty vector, wild-type, or designated deletion mutants were expressed in OK opossum kidney cells, the widely used model for PTH and FGF23 action on phosphate transport (13). FGF23 and PTH inhibited phosphate uptake in cells transfected with vector (Fig. 1C). Transfection with wild-type (WT) RGS14 abolished hormone-sensitive phosphate transport, as reported (5). Truncation mutants 1 and 2, where the N-terminus or N-terminus plus RGS domain, respectively, were eliminated (Fig. 1A), blocked FGF23 and PTH action, like full-length RGS14 (Fig. 1C), indicating that the critical regulatory regions lie further downstream. Indeed, removing the linker region with truncation construct 3, or the linker plus R1/R2 in construct 4 (Fig. 1A), abolished the physiological activity of RGS14 on FGF23- and PTH-regulated phosphate transport (Fig. 1C). These findings point to a vital function within the linker between the upstream RGS and downstream R1/R2 domains. Thus, the RGS14 PDZ ligand and an upstream component contained in the putative alpha-helical linker and not one of the signature RGS14 structural domains is responsible for the regulatory action of RGS14 on hormone-dependent phosphate transport.

### A Ser-rich sequence in the linker between RGS and R1 domains controls hormone action

We theorized that targeted phosphorylation of candidate Ser or Thr residues controls the response to hormone challenge. The linker between the RGS and R1 domains contains 18 Ser and 3 Thr residues. P.E.A.R.L, a post-translational modification prediction tool (14), identified a cluster of 5 Ser residues in the linker region as high-probability phosphorylation candidates (Fig. 2A). To test the hypothesis that Ser phosphorylation is required for RGS14 function, we initially generated an aggregate construct in which all 5 Ser residues were changed to Ala: Ser^260, 263, 266, 267, 269^Ala. Consistent with this hypothesis, RGS14-Ser^260, 263, 266, 267, 269^Ala abolished PTH- and FGF23 inhibition of phosphate uptake (Fig. 2B). To narrow down and determine if a single Ser residue within this regulatory sequence suffices to govern RGS14 functionality on PTH-sensitive phosphate transport, we generated individual Ser-to-Ala mutations at positions 260, 263, 266, 267, and 269. We tested and compared wild-type RGS14 with each discrete substitution to examine the effect on PTH inhibition of phosphate transport. In the absence of RGS14, PTH inhibited phosphate uptake (Fig. 2C). Wild-type RGS14 abolished PTH action, as did the Ser^260^Ala, Ser^263^Ala, and Ser^267^Ala constructs (Fig. 2C). In marked contrast, Ser^266^Ala and Ser^269^Ala failed to suppress RGS14 modulation of PTH inhibition of phosphate uptake (Fig. 2C). These results predict that Ser^266^Ala and Ser^269^Ala should not bind NHERF1 and, indeed, that was the case (Fig. 2D). Taken together, the findings are consistent with the view that Ser^266^ and Ser^269^ are critical sites for the actions of RGS14 and are compatible with a role for targeted phosphorylation in hormone-mediated inhibition of phosphate transport.

**Fig. 2.**
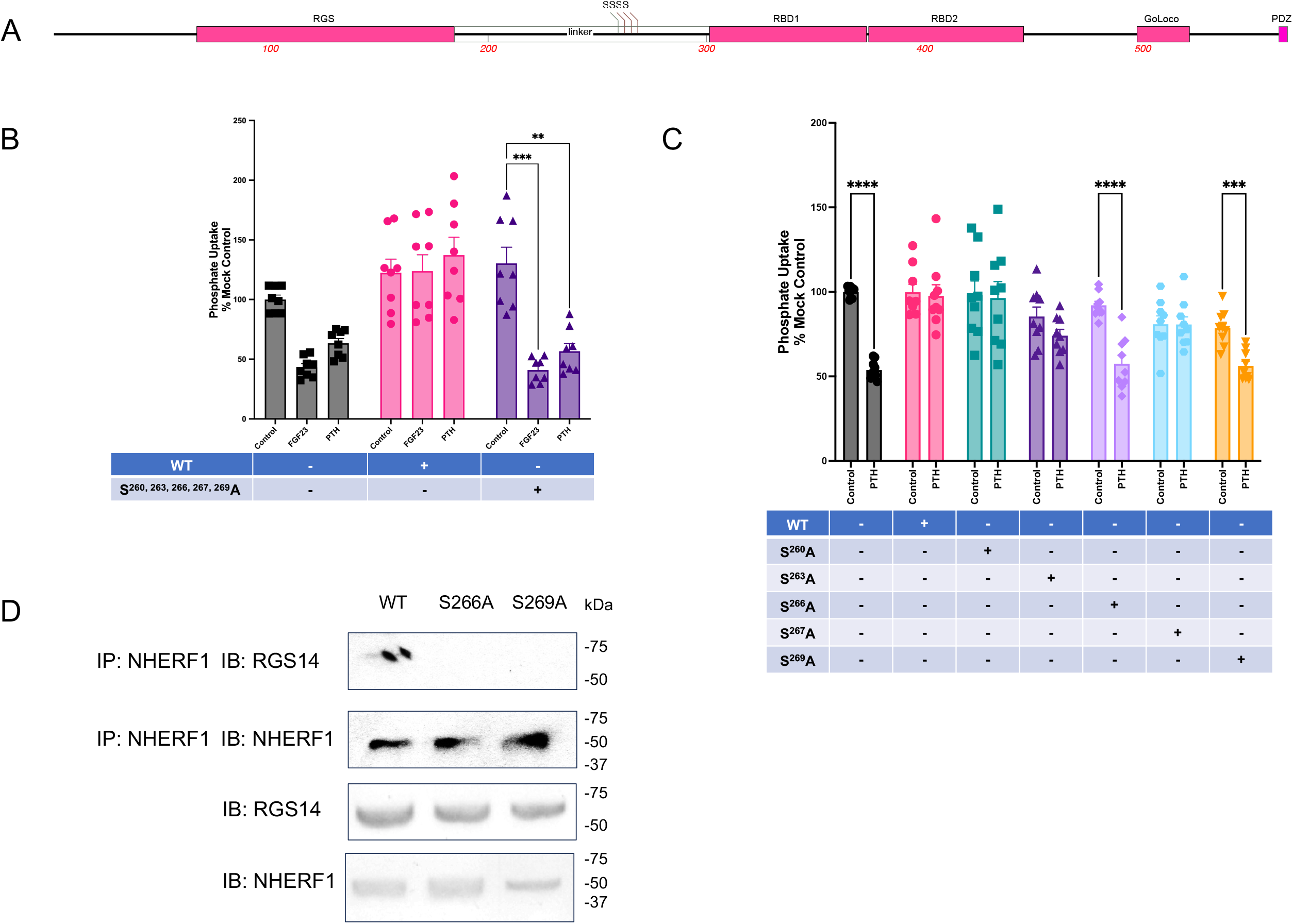
The linker sequence contains specific Ser residues that regulate RGS14 function. *A*. Schematic RGS14 sequence showing Ser locations in the linker region. *B*. Combined replacement of Ser with Ala at positions 260, 263, 266, 267, and 269 abolished the ability of RGS14 to regulate hormone inhibition of phosphate transport. OK cells were transfected with vector WT-RGS14, or ^S260, 263, 266, 267, 269^A-RGS14, as indicated. Hormone-sensitive phosphate uptake was measured and normalized to control conditions in vector-transfected cells (100%). *n*=4. ***p<0.0001 vs control. **p<0.001 vs. control. *C*. Substitution of individual Ser with Ala identifies residues in RGS14 controlling hormone-sensitive phosphate transport. OK cells were transfected with empty vector, WT-RGS14, or single Ser/Ala-RGS14 at positions 260, 263, 266, 267, and 269, as indicated. PTH-sensitive phosphate uptake was measured and normalized to control conditions in vector-transfected cells (100%). *n*=4. ***p<0.0005 vs control, ****p<0.0001 vs control. *D*. Ser^266^Ala and Ser^269^Ala-RGS14 do not immunoprecipitate with NHERF1. OK cells were transfected with HA-NHERF1 and FLAG-RGS14 (WT, Ser^266^Ala, Ser^269^Ala). 48 hr after transfection, lysates were prepared and NHERF1 was immunoprecipitated using monoclonal anti-HA agarose (Sigma). RGS14 was immunoblotted with a rabbit polyclonal anti-FLAG antibody (Sigma). NHERF1 was immunoblotted using a rabbit polyclonal anti-HA antibody (Sigma). Molecular weight markers are depicted on the right of the blots (kDa). *n*=1-2.

## Discussion

Twenty canonical mammalian RGS proteins have been described and grouped into four families based on sequence homologies and the presence or absence of included domains (15). Their distinguishing action, mediated by their shared RGS domains, accelerates GTPase activity by hydrolyzing bound GTP, thereby terminating G protein signaling. RGS14 is a family D (15) or R12 (16) member that includes RGS12 and RGS10. The biological functions of RGS proteins are incompletely understood but include tonic suppression of synaptic plasticity and hippocampal-based learning (8,17,18), reduced myocardial remodeling (19), and altered adipose tissue metabolism (20). A pioneering GWAS study linked *RGS14* variants to differences in circulating PTH levels (21). Although the identified SNP was intronic, other coding variants in the RGS14 PDZ ligand (22) prompted us to explore whether it bound NHERF1 and how this affected phosphate uptake. Naturally occurring RGS14 Asp^563^Asn, for instance, disrupted binding to NHERF1 and, predictably, blocked PTH-sensitive Pi transport (5). By contrast, the Asp^563^Gly variant bound NHERF1 and supported PTH-inhibitable Pi uptake. PDZ ligand residues are numbered backward from zero at the carboxy terminus. The -1 position is considered permissive and, indeed, Ala^565^ variants at this site were fully tolerated and behaved like wild-type RGS14, abolishing PTH inhibition of Pi uptake. Notably, mutations that interfered with RGS14 binding to NHERF1 also hampered RGS14 capacity to affect PTH-regulated Pi transport. The data highlighted a close correlation between RGS14 binding to NHERF1 and its ability to restrict PTH-sensitive Pi transport.

RGS14 strikingly governed both PTH and FGF23 actions on phosphate transport. Its cognate GPCR receptor mediates PTH effects, while FGFR1 transmits FGF23 effects on phosphate transport (4), a receptor tyrosine kinase unrelated to transmembrane GPCRs. This observation made it unlikely that RGS14 regulatory activity stemmed from G protein activity or one of the integral RGS14 domains associated with G protein function. Accordingly, we initiated the current studies to delineate the RGS14 component responsible for hormone-regulated actions. We employed a strategy involving sequential elimination of the defined regions of RGS14. The results (Fig. 1C) disclosed that none of the signature RGS14 domains was responsible for the phosphate transport regulatory activity. Instead, the findings revealed participation of the linker region between the RGS and R1 domains.

We postulated that targeted phosphorylation of one or more of the 21 Ser and Thr residues in the linker was the required key to unlocking RGS14 regulatory actions. Software analysis of the region distinguished 5 high-value sites that we initially focused on. All 5 identified Ser were replaced with Ala for an initial analysis. This block approach supported the view that regulation could be attributed to this subset of polar residues. Further examination with single Ser/Ala exchange replacements assigned critical roles to Ser^266^ and Ser^269^, which behaved the same as wild-type RGS14 (Fig. 2C). These findings do not exclude the participation of additional Ser or Thr residues within the linker. They also do not speak to possible sequential phosphorylation events. The PTHR and FGFR1 signaling pathways impinging on RGS14 remain to be defined. Earlier work disclosed that PTH and FGF23 exert shared actions on phosphate excretion in some instances, while their effects are singular and discrete in others (4). PTH and FGF23 share select phosphorylation sites on NHERF1 (23). Whether or not a mutually activated kinase is responsible for Ser phosphorylation of the RGS14 linker is unknown. The alpha-helical interlinker contains a dibasic PKA consensus sequence (RRXS/T) that includes Ser^260^. However, this residue did not participate in ablating PTH-sensitive phosphate transport (Fig. 2C). Neither Ser^266^ nor Ser^269^, which eliminated RGS14 activity, restoring PTH regulation, are part of a described conserved or degenerate consensus linear phosphorylation motif. Nonetheless, <u>NetPhos 3.1</u> predicts phosphorylation with a 0.99 probability of the sequence harboring Ser^266^ and Ser^269^ (24). Recent work suggests that kinases may recognize secondary structure (25), and this may account for the atypical behavior

RGS14 is reported to be highly phosphorylated (22), and considerable evidence indicates that its phosphorylation state regulates its actions. Rat Rgs14 is phosphorylated by PKA at Thr^494^ outside the linker region to regulate Gαi1-GDP interactions (26). Within the linker region, Rgs14 is phosphorylated at Ser^218^ to govern 14-3-3γ binding and nuclear localization (27). Consistent with these findings, we show here that phosphorylation of RGS14 at either Ser^266^ or Ser^269^ regulates its actions towards hormone-sensitive phosphate uptake in kidney cells. Additional work beyond the scope of the present investigation will be needed to define the kinases and phosphatases responsible for Ser phosphocycling and search for other candidate Ser or Thr residues involved.

We conclude that the RGS14 linker and its PDZ ligand are critical for regulating control of PTH- and FGF23-sensitive phosphate transported mediated by NPT2A. Earlier studies disclosed that RGS14 binds NHERF1 PDZ2 (5).

## Experimental procedures

### Chemical reagents, plasmids, and antibodies

Monoclonal anti-HA agarose (A2095) and rabbit polyclonal anti-FLAG (F7425) were obtained from Sigma. Rabbit polyclonal anti-HA (sc-805) and protein G+ agarose (sc-2002) were purchased from Santa Cruz Biotechnology. Mouse and rabbit polyclonal anti-NHERF1 antibodies were purchased from Abcam (ab-9526 and ab3452, respectively). Rabbit polyclonal anti-RGS14 was purchased from Proteintech (16258-1-AP). We purchased anti-human-NPT2A from Novus Biologicals (NBP242216) and anti-actin from Santa Cruz Biotechnology (sc-1616R). HA-NHERF1 as described (28). [Nle^8,18^,Tyr^34^]PTH(1–34) was from Bachem, Torrance, CA (H9110). Recombinant human Arg^179^Gln-FGF23^25-251^ (herein referred to as FGF23), which is resistant to furin cleavage and inactivation, was obtained from R&D Systems (2604-FG-025).

### Human RGS14 constructs

FLAG-tagged deletion mutants of RGS14 were created using the full-length human RGS14-pcDNA3.1 as a template for PCR using primers listed in Table 1. All PCR products were digested and cloned into pcDNA3.1+ between the HindIII and EcoRI restriction sites.

**Table 1.**
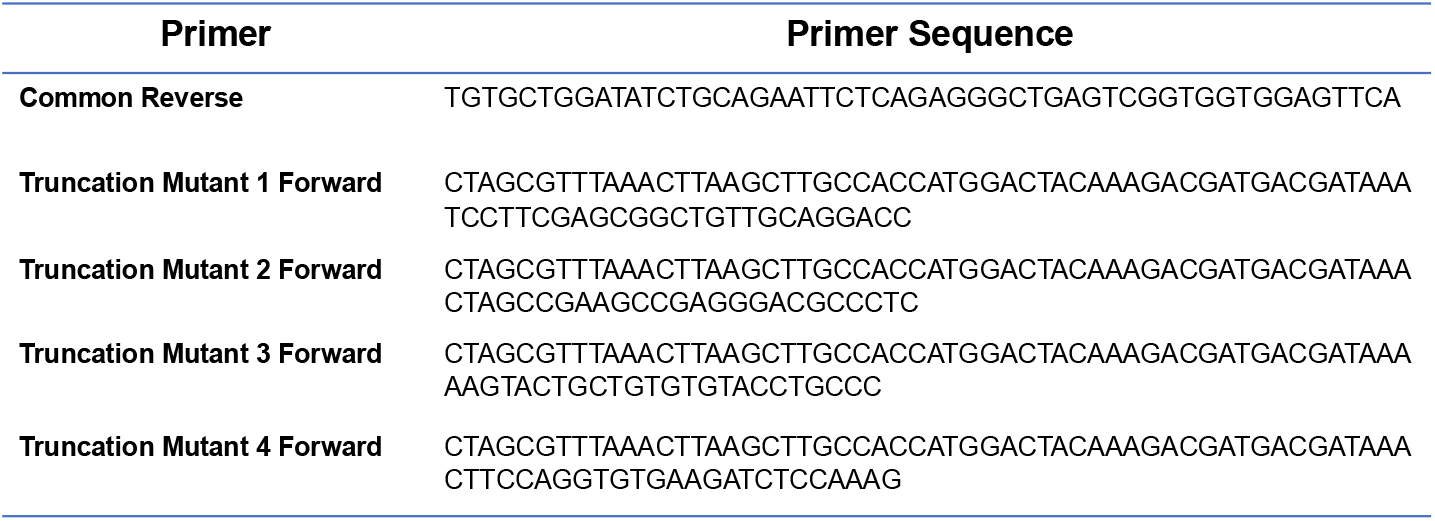
Primers used to generate RGS14 truncation mutants.

Single Ser-to-Ala substitutions were introduced into FLAG-RGS14 using the Agilent QuikChange Lightning Site-Directed Mutagenesis Kit (210519) and primers detailed in Table 2.

**Table 2.**
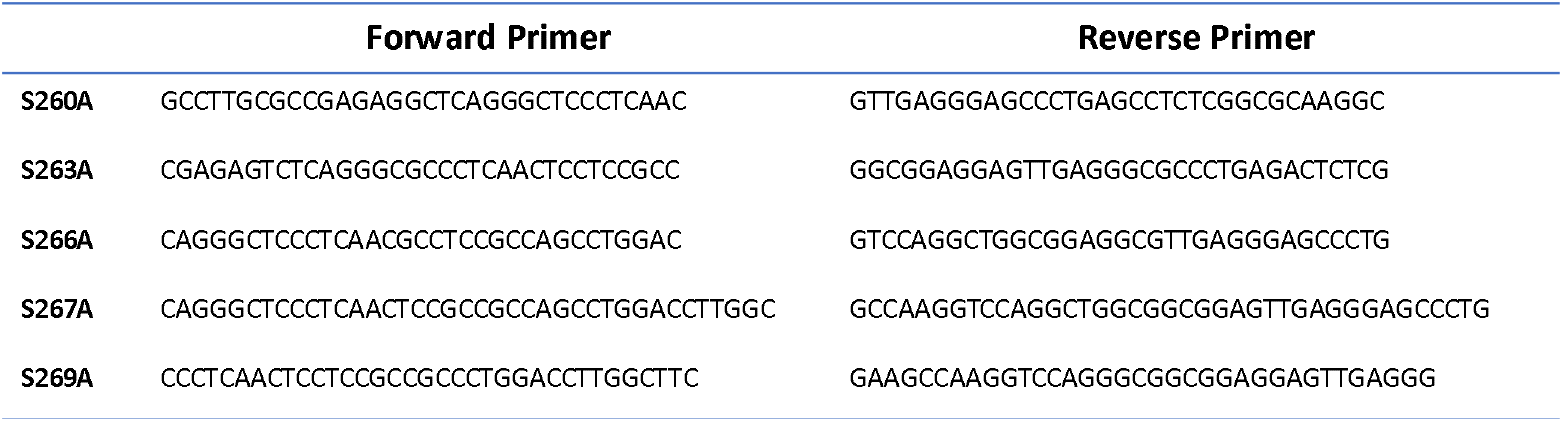
Primers used to generate single RGS14 Ser-Ala mutations in the linker region.

### Cell lines and culture

Opossum kidney cells (OK/B) were obtained from J. Cole (29) and cultured in DMEM/F-12 (Corning, 10-090-CV) supplemented with 5% heat-inactivated fetal bovine serum (GenClone 25-514H, Genesee Scientific) plus 1% penicillin and streptomycin.

Human Embryonic Kidney (HEK293) cells were cultured in DMEM/F12 (Corning, 10-090-CV) supplemented with 10% heat-inactivated fetal bovine serum (GenClone 25-514H, Genesee Scientific) plus 1% penicillin and streptomycin.

### Pi transport

OK cells were seeded on 12-well plates. 24 hr later, as indicated, cells were transfected with 1μg/well of WT or mutant FLAG-RGS14 plasmid, using Lipofectamine 3000 (Invitrogen). After 48 hr, cells were serum-starved overnight and then treated for 2 hr with 100 nM PTH(1-34) or FGF23, as indicated. Pi uptake was measured as described (4). The hormone-supplemented medium was aspirated, and the wells were washed three times with 1 ml of Na-replete wash buffer (140 mM NaCl, 4.8 mM KCl, 1.2 mM MgSO_4_, 0.1 mM KH_2_PO_4_, 10 mM HEPES, pH 7.4). The cells were incubated with 1 μCi [^32^P]orthophosphate (PerkinElmer Life Sciences, NEX053) in 1 ml of Na-replete buffer for 10 min. Pi uptake was terminated by placing the plate on ice and rinsing the cells three times with Na-free buffer (140 mM N-methyl-D-glucamine, 4.8 mM KCl, 1.2 mM MgSO_4_, 0.1 mM KH_2_PO_4_, 10 mM HEPES, pH 7.4). The cells in each well were extracted overnight at 4 °C using 500 μl 1% Triton X-100 (Sigma). A 250-μl aliquot was counted in a Beckmann Coulter LS6500 scintillation spectrometer. Data were normalized to Pi uptake under control conditions defined as 100%.

### Transfection and Immunoblotting

As indicated, protein lysates were prepared from OK and HEK293 cells transfected with various RGS14. Cells were seeded on 6-well plates. 24 hr later, the cells were transfected with 1μg/well of human RGS14 expression plasmid using Lipofectamine 3000 (Invitrogen). After 48 hr, protein lysates were prepared using 1% Nonidet P-40 buffer supplemented with protease inhibitor mixture I.

Cells (OK/B, HEK293) were seeded on 6-well plates concurrently with the 12 well plates used in the phosphate uptake measurements. After 24 hr, cells were transfected with 1μg each/well of HA-NHERF1 and/or FLAG-RGS14 (WT, mutants as indicated) using Lipofectamine 3000 (Invitrogen). 48 hr post-transfection, cells were lysed with 1% Nonidet P-40 (50 mM Tris, 150 mM NaCl, 5 mM EDTA, 1% Nonidet P-40) supplemented with protease inhibitor mixture I (EMD Millipore). Lysis was performed for 15 min on ice, and membrane components were pelleted at 12000 rpm for 15 min at 4 °C in an Eppendorf 5415-R refrigerated microcentrifuge. The supernatant was mixed with an equal volume of SDS-PAGE sample buffer (Bio-Rad) supplemented with 5% β-mercaptoethanol (Sigma). These protein samples were resolved on 10% SDS-polyacrylamide gels and transferred to Immobilon-P membranes (Millipore) using the semidry method (Bio-Rad). Membranes were blocked for 1 hr at room temperature with 5% nonfat dried milk in Tris-buffered saline plus Tween 20 (TBST) (blocking buffer) and incubated with the indicated primary antibodies at a dilution of 1:1000 in blocking buffer overnight at 4 °C. The membranes were washed four times for 10 min in TBST and then incubated with the secondary antibodies (goat anti-rabbit IgG or donkey anti-mouse IgG conjugated to horseradish peroxidase) at a 1:5000 dilution for 1 hr at room temperature. Membranes were washed four times for 10 min in TBST. Protein bands were detected by Luminol-based enhanced chemiluminescence (EMD Millipore WBKLS0500). Immunoblots were captured on film and quantified using Image J.

### Statistical analysis

Results were analyzed using Prism 10 software (GraphPad, La Jolla, CA). Data represent the mean ± SD or SEM as indicated of *n* ≥ 3 independent experiments and were compared by analysis of variance with *post hoc* testing using the Bonferonni procedure or paired t-test as appropriate. *p* values < 0.05 were considered statistically significant.

## Footnotes

^2^Human proteins are indicated by 3-letter uppercase convention; genes in italics. Only the first letter is uppercase for the corresponding mouse protein or gene.

## The abbreviations used are

PTH: parathyroid hormone
PTHR: PTH receptor
NPT2A: Sodium-phosphate cotransport protein 2A
RGS14: regulator of G protein signaling 14
PDZ: postsynaptic density protein 95 (PSD95), drosophila disc large tumor suppressor (DlgA), and zonula occludens 1 protein (ZO1)
GAP: GTPase activating protein
RBD: Ras-binding domain
GPR: G protein regulatory motif
HEK293: Human Embryonic Kidney 293 cells.

## Findings and additional information

This work was supported by NIH grant GM140632-A1 (JRH, PAF).

## Footnotes

2 human RGS14 protein is denoted by upper case letters; only the first letter is capitalized for other species.

## References

1. Mahon, M. J., Donowitz, M., Yun, C. C., and Segre, G. V. (2002) Na+/H+ exchanger regulatory factor 2 directs parathyroid hormone 1 receptor signalling. Nature 14, 858–861

2. Sneddon, W. B., Syme, C. A., Bisello, A., Magyar, C. E., Weinman, E. J., Rochdi, M. D., Parent, J. L., Abou-Samra, A. B., and Friedman, P. A. (2003) Activation-independent parathyroid hormone receptor internalization is regulated by NHERF1 (EBP50). J. Biol. Chem. 14, 43787–43796

3. Weinman, E. J., Steplock, D., Shenolikar, S., and Biswas, R. (2011) Fibroblast growth factor-23-mediated inhibition of renal phosphate transport in mice requires sodium-hydrogen exchanger regulatory factor-1 (NHERF-1) and synergizes with parathyroid hormone. J. Biol. Chem. 14, 37216–37221

4. Sneddon, W. B., Ruiz, G. W., Gallo, L. I., Xiao, K., Zhang, Q., Rbaibi, Y., Weisz, O. A., Apodaca, G. L., and Friedman, P. A. (2016) Convergent signaling pathways regulate parathyroid hormone and fibroblast growth factor-23 action on NPT2A-mediated phosphate transport. J. Biol. Chem. 14, 18632–18642

5. Friedman, P. A., Sneddon, W. B., Mamonova, T., Montanez-Miranda, C., Ramineni, S., Harbin, N. H., Squires, K. E., Gefter, J. V., Magyar, C. E., Emlet, D. R., and Hepler, J. R. (2022) RGS14 regulates PTH-and FGF23-sensitive NPT2A-mediated renal phosphate uptake via binding to the NHERF1 scaffolding protein. J. Biol. Chem. 14, 101836

6. Almutairi, F., Lee, J. K., and Rada, B. (2020) Regulator of G protein signaling 10: Structure, expression and functions in cellular physiology and diseases. Cell. Signal. 14, 109765

7. Harbin, N. H., Bramlett, S. N., Montanez-Miranda, C., Terzioglu, G., and Hepler, J. R. (2021) RGS14 Regulation of Postsynaptic Signaling and Spine Plasticity in Brain. Int. J. Mol. Sci. 22

8. Evans, P. R., Gerber, K. J., Dammer, E. B., Duong, D. M., Goswami, D., Lustberg, D. J., Zou, J., Yang, J. J., Dudek, S. M., Griffin, P. R., Seyfried, N. T., and Hepler, J. R. (2018) Interactome analysis reveals regulator of G Protein signaling 14 (RGS14) is a novel calcium/calmodulin (Ca2+/CaM) and CaM Kinase II (CaMKII) binding partner. J. Proteome Res. 14, 1700–1711

9. Shu, F. J., Ramineni, S., and Hepler, J. R. (2010) RGS14 is a multifunctional scaffold that integrates G protein and Ras/Raf MAPkinase signalling pathways. Cell. Signal. 14, 366–376

10. Brown, N. E., Goswami, D., Branch, M. R., Ramineni, S., Ortlund, E. A., Griffin, P. R., and Hepler, J. R. (2015) Integration of G protein α (Gα) signaling by the regulator of G protein signaling 14 (RGS14). J. Biol. Chem. 14, 9037–9049

11. Snow, B. E., Antonio, L., Suggs, S., Gutstein, H. B., and Siderovski, D. P. (1997) Molecular cloning and expression analysis of rat Rgs12 and Rgs14. Biochem. Biophys. Res. Commun. 14, 770–777

12. Vellano, C. P., Brown, N. E., Blumer, J. B., and Hepler, J. R. (2013) Assembly and function of the regulator of G protein signaling 14 (RGS14).H-Ras signaling complex in live cells are regulated by Galphai1 and Galphai-linked G protein-coupled receptors. J. Biol. Chem. 14, 3620–3631

13. Biber, J., Malmström, K., Reshkin, S., and Murer, H. (1990) Phosphate transport in established renal epithelial cell lines. Methods Enzymol. 14, 494–504

14. English, N., and Torres, M. (2022) Enhancing the Discovery of Functional Post-Translational Modification Sites with Machine Learning Models - Development, Validation, and Interpretation. Methods Mol. Biol. 14, 221–260

15. Stewart, A., and Fisher, R. A. (2015) Introduction: G Protein-coupled receptors and RGS proteins. Prog. Mol. Biol. Transl. Sci. 14, 1–11

16. Ross, E. M., and Wilkie, T. M. (2000) GTPase-activating proteins for heterotrimeric G proteins: regulators of G protein signaling (RGS) and RGS-like proteins. Annu. Rev. Biochem. 14, 795–827

17. Evans, P. R., Dudek, S. M., and Hepler, J. R. (2015) Regulator of G Protein Signaling 14: A Molecular Brake on Synaptic Plasticity Linked to Learning and Memory. Prog. Mol. Biol. Transl. Sci. 14, 169–206

18. Lee, S. E., Simons, S. B., Heldt, S. A., Zhao, M., Schroeder, J. P., Vellano, C. P., Cowan, D. P., Ramineni, S., Yates, C. K., Feng, Y., Smith, Y., Sweatt, J. D., Weinshenker, D., Ressler, K. J., Dudek, S. M., and Hepler, J. R. (2010) RGS14 is a natural suppressor of both synaptic plasticity in CA2 neurons and hippocampal-based learning and memory. Proc. Natl. Acad. Sci. U. S. A. 14, 16994–16998

19. Li, Y., Tang, X. H., Li, X. H., Dai, H. J., Miao, R. J., Cai, J. J., Huang, Z. J., Chen, A. F., Xing, X. W., Lu, Y., and Yuan, H. (2016) Regulator of G protein signalling 14 attenuates cardiac remodelling through the MEK-ERK1/2 signalling pathway. Basic Res. Cardiol. 14, 47

20. Vatner, D. E., Zhang, J., Oydanich, M., Guers, J., Katsyuba, E., Yan, L., Sinclair, D., Auwerx, J., and Vatner, S. F. (2018) Enhanced longevity and metabolism by brown adipose tissue with disruption of the regulator of G protein signaling 14. Aging Cell 14, e12751

21. Robinson-Cohen, C., Lutsey, P. L., Kleber, M. E., Nielson, C. M., Mitchell, B. D., Bis, J. C., Eny, K. M., Portas, L., Eriksson, J., Lorentzon, M., Koller, D. L., Milaneschi, Y., Teumer, A., Pilz, S., Nethander, M., et al. (2017) Genetic variants associated with circulating parathyroid hormone. J. Am. Soc. Nephrol. 14, 1553–1565

22. Squires, K. E., Montanez-Miranda, C., Pandya, R. R., Torres, M. P., and Hepler, J. R. (2018) Genetic analysis of rare human variants of regulators of G protein signaling proteins and their role in human physiology and disease. Pharmacol. Rev. 14, 446–474

23. Zhang, Q., Xiao, K., Paredes, J. M., Mamonova, T., Sneddon, W. B., Liu, H., Wang, D., Li, S., McGarvey, J. C., Uehling, D., Al-awar, R., Joseph, B., Jean-Alphonse, F., Orte, A., and Friedman, P. A. (2019) Parathyroid hormone initiates dynamic NHERF1 phosphorylation cycling and conformational changes that regulate NPT2A-dependent phosphate transport. J. Biol. Chem. 14, 4546–4571

24. Blom, N., Sicheritz-Ponten, T., Gupta, R., Gammeltoft, S., and Brunak, S. (2004) Prediction of post-translational glycosylation and phosphorylation of proteins from the amino acid sequence. Proteomics 14, 1633–1649

25. Miller, C. J., and Turk, B. E. (2018) Homing in: Mechanisms of substrate targeting by protein kinases. Trends Biochem. Sci. 14, 380–394

26. Hollinger, S., Ramineni, S., and Hepler, J. R. (2003) Phosphorylation of RGS14 by protein kinase A potentiates its activity toward Gαi. Biochemistry 14, 811–819

27. Gerber, K. J., Squires, K. E., and Hepler, J. R. (2018) 14-3-3γ binds regulator of G protein signaling 14 (RGS14) at distinct sites to inhibit the RGS14:Gαi-AlF4-signaling complex and RGS14 nuclear localization. J. Biol. Chem. 14, 14616–14631

28. Wang, B., Means, C. K., Yang, Y., Mamonova, T., Bisello, A., Altschuler, D. L., Scott, J. D., and Friedman, P. A. (2012) Ezrin-anchored PKA coordinates phosphorylation-dependent disassembly of a NHERF1 ternary complex to regulate hormone-sensitive phosphate transport. J. Biol. Chem. 14, 24148–24163

29. Cole, J. A., Forte, L. R., Krause, W. J., and Thorne, P. K. (1989) Clonal sublines that are morphologically and functionally distinct from parental OK cells. Am. J. Physiol. 14, F672–679

